# Glycoproteomic measurement of site-specific polysialylation

**DOI:** 10.1101/740928

**Authors:** Ruby Pelingon, Cassandra L. Pegg, Lucia F. Zacchi, Toan K. Phung, Christopher B. Howard, Ping Xu, Matthew P. Hardy, Catherine M. Owczarek, Benjamin L. Schulz

## Abstract

Polysialylation is the enzymatic addition of a highly negatively charged sialic acid polymer to the non-reducing termini of glycans. Polysialylation plays an important role in development, and is involved in neurological diseases, neural tissue regeneration, and cancer. Polysialic acid (PSA) is also a biodegradable and non-immunogenic conjugate to therapeutic drugs to improve their pharmacokinetics. PSA chains vary in length, composition, and linkages, while the specific sites of polysialylation are important determinants of protein function. However, PSA is difficult to analyse by mass spectrometry (MS) due to its high negative charge and size. Most analytical approaches for analysis of PSA measure its degree of polymerization and monosaccharide composition, but do not address the key questions of site specificity and occupancy. Here, we developed a high-throughput LC-ESI-MS/MS glycoproteomics method to measure site-specific polysialylation of glycoproteins. This method measures site-specific PSA modification by using mild acid hydrolysis to eliminate PSA and sialic acids while leaving the glycan backbone intact, together with protease digestion followed by LC-ESI-MS/MS glycopeptide detection. PSA-modified glycopeptides are not detectable by LC-ESI-MS/MS, but become detectable after desialylation, allowing measurement of site-specific PSA occupancy. This method is an efficient analytical workflow for the study of glycoprotein polysialylation in biological and therapeutic settings.

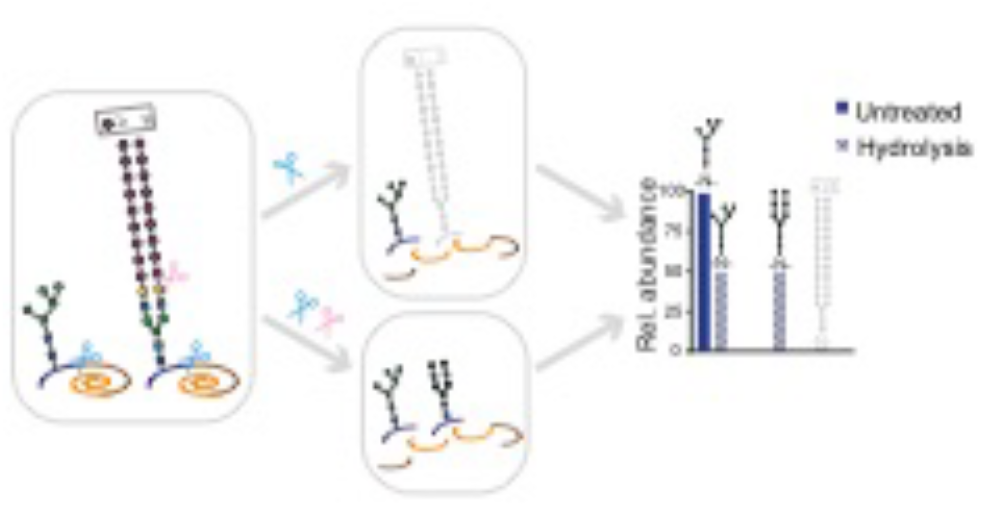
Graphical Abstract.

## Introduction

Protein glycosylation, the enzymatic addition of carbohydrates to a protein, is a common post-translational modification that plays a variety of physiological roles.^1^ Protein glycosylation is critical in protein folding and transport through the secretory pathway, and is also important for protein solubility and stability, cell-cell interactions, and ligand-receptor binding.^1–4^ *N*-linked glycosylation occurs at selected asparagine (N) residues within the consensus “sequon” N-X-S/T (where X ≠ proline).^5^ *N*-linked glycans are pre-assembled on the membrane of the endoplasmic reticulum (ER) as a dolichol-linked Glc(3)Man(9)GlcNAc(2) structure (Glc, glucose; Man, mannose; and GlcNAc, *N*-Acetylglucosamine), and are transferred *en bloc* by the enzyme oligosaccharyltransferase to the polypeptide in the lumen of the ER.^3^ Glycans are modified as proteins traverse the secretory pathway, leading to heterogeneity in the glycan structures in mature glycoproteins.^1, 6, 7^ The presence and structures of glycans at particular sites have a major influence on protein conformation, stability, and function.^8^

Mammalian glycans are commonly terminated by sialic acids, 9-carbon carboxylated monosaccharides.^9, 10^ Although there are many sialic acid variants, N-acetylneuraminic acid (NeuAc) is the most common in human glycans. Glycans are typically monosialylated, but sialic acids can also be found as short oligosialyl chains (2-7 monosaccharides) and as polysialic acid (PSA) (≥ 8 monosaccharides), with an arbitrary length distinction between oligo- and polysialic acid.^9, 11^ PSA is structurally heterogenous due to variability in the sialic acid type, modifications, glycosidic linkages, and degree of polymerization.^10^ Only few polysialylated proteins have been identified, including neural cell adhesion molecule (NCAM),^12^ synCAM-1,^13^ neuropilin-2,^14^ voltage-sensitive sodium channel,^15, 16^ CCR7 chemokine,^17^ E-selecting ligand-1,^18^ CD36 in milk,^19^ and the polysialyltransferases themselves.^20, 21^ NCAM is the best-studied polysialylated protein, and carries α2-8-linked NeuAc PSA in two of its *N*-glycans.^11, 22–24^ Bulky and negatively charged PSA-modified NCAM inhibits cell-cell interactions, thereby playing important roles in embryonic development,^11^ neurological diseases,^11, 25, 26^ tumour metastasis,^27^ and neural tissue regeneration.^28^ PSA also has important biotechnological applications.^11, 29^ PSA is used as a protein conjugate (chemically/enzymatically attached to proteins) or as a delivery system (PSA micelles) to improve pharmacokinetics of therapeutic proteins.^29–34^ Thus, polysialylation is a post-translational modification with high physiological, pathological, and biotechnological relevance.

Analysis of PSA-modified glycoproteins is challenging due to the large size, high negative charge, and structural heterogeneity of PSA. Immuno-detection can detect protein polysialylation,^10, 35, 36^ and detection with and without endosialidase treatment allows inference of the presence of PSA-modified proteins.^37–39^ However, anti-PSA immunodetection provides no information about the sites of polysialylation, PSA structure and composition, the degree of polymerization, or the underlying glycan structures. Several approaches overcome some of these limitations.^40, 41^ PSA degree of polymerization, sialic acid composition, and glycosidic linkages can be measured using: mild acid hydrolysis, derivatization, and HPLC;^42–45^ fluorometric C7/C9 analysis;^40, 45–47^ high performance (high pH) anion-exchange chromatography pulsed electrochemical detection (HPAEC-PED);^48, 49^ nuclear magnetic resonance (NMR);^50–53^ or mass spectrometry (MS) glycomics ^54–57^, including matrix-assisted laser desorption/ionization time-of-flight MS (MALDI-TOF MS),^56, 58, 59^ gas chromatography-MS (GC-MS),^60^ and electrospray ionization MS (ESI-MS).^57, 61^ Therefore, a wide range of chemical and analytical tools are available to characterize released PSA chains. Three additional features of polysialylated glycoproteins are critical: site-specific PSA localisation, PSA occupancy, and the underlying structure of the polysialylated glycan. Sites of PSA modification can be identified by site-directed mutagenesis of glycosylation sites with anti-PSA Western blotting,^22, 62^ or immunoenrichment of tryptic PSA-glycopeptides, followed by deglycosylation and MS analysis.^13, 51, 63, 64^ However, these methods do not measure site-specific PSA occupancy.

Here, we developed a high-throughput glycoproteomic workflow to measure site-specific polysialylation of glycoproteins (Figure 1). This workflow combines mild acid hydrolysis to remove PSA from glycans, followed by LC-ESI-MS/MS for detection and site-specific relative quantification of previously polysialylated asialoglycopeptides (Figure 1). By comparing glycoform abundance in the desialylated sample with that in the untreated control, site specific polysialylation can be inferred, its occupancy measured, and the underlying polysialylated glycan structure can be elucidated.

**Figure 1.**
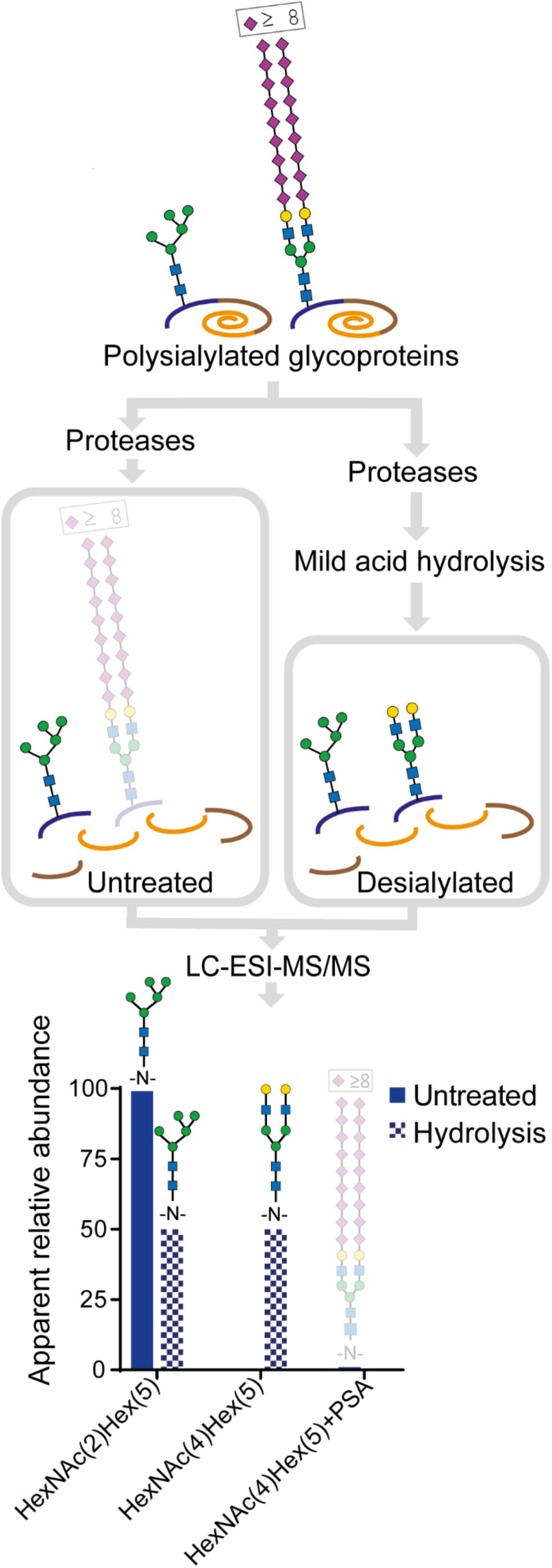
Schematic representation of our glycoproteomic method to detect and measure glycoprotein polysialylation. Polysialylated glycoproteins are protease digested to peptides and analysed by LC-ESI-MS/MS with and without desialylation by mild acid hydrolysis. Comparison of apparent site-specific glycoform abundance after desialylation and in untreated controls allows inference of site-specific glycan polysialylation and occupancy.

## Experimental Section

### Cloning, expression, and purification of polysialylated recombinant human NCAM

cDNAs encoding human Neural Cell Adhesion Molecule 1 (NCAM) (GenBank Accession number NP_000054), human β1,4-galactosyltransferase 1 (B4GALT1) (NP_001497), human CMP-*N*-acetylneuraminate-β-galactosamide-α-2,3-sialyltransferase 4 (ST3GAL4) (NP_001241686), and human CMP-*N*-acetylneuraminate-poly-α-2,8-sialyltransferase (ST8SIA4) (NP_005659)were codon-optimized for human expression and synthesized (Geneart, Thermo Fisher Scientific). A cDNA encoding a soluble version of human NCAM, with a stop codon introduced after amino acid 718, was generated using standard PCR-based techniques, with a C-terminal 8His tag and a FLAG (DYKDDDDK) tag after the signal peptide cleavage site (rHuNCAM, Figure 2 and Supplementary Figure 2). cDNAs had a Kozak consensus sequence (GCCACC) placed upstream of the initiating methionine. PCR products were digested with NheI and XhoI, and ligated into pcDNA3.1 (Invitrogen, Thermo Fisher Scientific) to produce pcDNA-HuNCAM(1-19)-FLAG-HuNCAM(20-718)-8His. Plasmid DNA was prepared using QIAGEN Plasmid Giga Kits according to manufacturer’s instructions.

**Figure 2.**
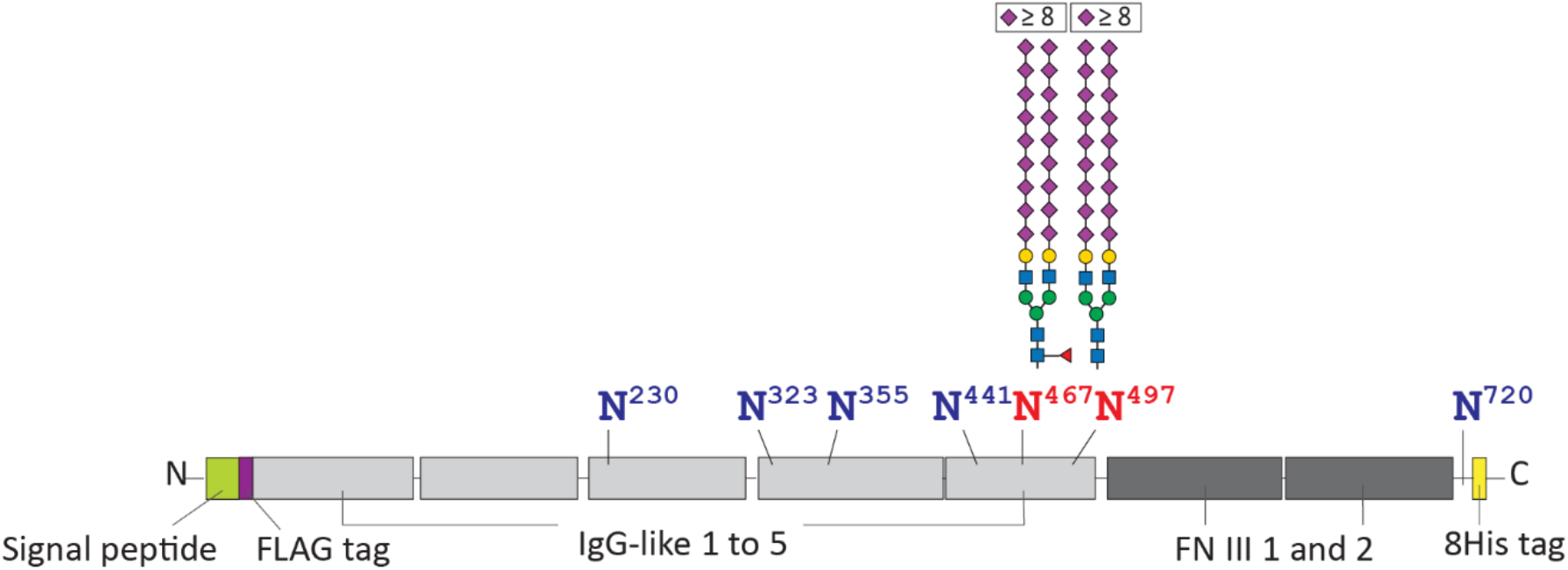
*N*-linked glycosylation sites on rHuNCAM. The protein domain structure of rHuNCAM is shown as per UniProtKB (P13591), with five Ig-like domains (light grey) and two fibronectin type III (FN) domains (dark grey), joined by short linker regions of variable size. The protein is truncated at amino acid 718 to generate a soluble version of HuNCAM; a FLAG tag (violet) is located at the N-terminus after the signal sequence (light green); and an 8His tag is attached to the C-terminal end of rHuNCAM (yellow). There are six *N*-linked glycosylation sites in NCAM’s Ig-like domains and one *N*-linked site close to the transmembrane domain. The 5^th^ and 6^th^ *N*-linked sites at N467 and N496 of rHuNCAM are reported to be polysialylated in NCAM (N459 and N488, as per native human protein sequence).^11, 22–24^

Expi293F cells (Invitrogen, Thermo Fisher Scientific) were cultured in Expi293 Expression Medium (Invitrogen, Thermo Fisher Scientific) supplemented with Antibiotic-Antimycotic (GIBCO, Thermo Fisher Scientific) and cells were maintained in an incubator at 37 °C and 150 rpm with an atmosphere of 8 % CO_2_. Transient transfections of expression plasmids used the Expi293 Expression system and were performed following manufacturer’s recommendations. To produce polysialylated rHuNCAM, expression plasmids pcDNA-HuNCAM(1-19)-FLAG-HuNCAM(20-718)-8His, pcDNA-Hu-ST8SIA4, pcDNA-Hu-B4GALT1, and pcDNA-Hu-ST3GAL4 with DNA ratios of 0.85:0.05:0.05:0.05 were co-transfected into Expi293 cells, with 0.5 % (w/v) LucraTone Lupin (Celliance) added 18-20 h post-transfection.^65, 66^ After 6 days of incubation cells were harvested as described.^67^ Expression of secreted polysialylated rHuNCAM was verified by SDS-PAGE and Coomassie Blue staining, and Western blot analysis using anti-His (A01620, Genscript) and anti-PSA-NCAM (MAB5324, Millipore) antibodies. rHuNCAM was purified from the culture supernatant of transiently transfected Expi293F cells using nickel sepharose resin and size-exclusion chromatography as previously described.^68, 69^

### Expression, purification, and activity of recombinant Endosialidase NF

The pCW(-lacZ) vector containing the MalE-thrombin-Endosialidase NF construct was a kind gift from Warren Wakarchuk.^37^ Expression and purification of MalE-thrombin-Endosialidase NF (EndoNF) used a published method with minor modifications.^70^ EndoNF retained in the amylose-agarose resin was eluted with working buffer supplemented with 10 mM maltose. The purity and quality of EndoNF were determined by SDS-PAGE and Coomassie Blue staining (Supplementary Figure 1).

To enzymatically de-polysialylate rHuNCAM, 5 μg of rHuNCAM was diluted in 50 mM sodium phosphate buffer pH 7.5, treated or not with 0.25 μg EndoNF (20:1 sample:EndoNF ratio), and incubated at 37 °C and 1500 rpm for 2 h in a MS100 thermoshaker incubator (LabGene Scientific). Samples were precipitated with 4 volumes of 1:1 acetone:methanol, incubated at −20 °C for 16 h, and centrifuged at 20,000 rcf for 20 min at 4 °C. Pellets were reconstituted in 1X SDS-PAGE loading buffer, incubated at 65 °C for 15 min, and separated by SDS-PAGE using a 6% polyacrylamide gel. Separated proteins were either stained with Coomassie Blue or transferred onto nitrocellulose membrane (Bio-Rad) using an XCell II Blot Module (Invitrogen, Thermo Fisher Scientific). Western blots using anti-PSA antibody (mAb735, Abcam) were developed using SuperSignal West Dura Extended Duration Substrate (Thermo Fisher Scientific), and images were captured using a GE Amersham Imager 600.

### Mass Spectrometry sample preparation

Four treatments were tested: EndoNF alone, EndoNF followed by mild acid hydrolysis, mild acid hydrolysis only, and untreated control. For each treatment, 5 μg of purified rHuNCAM or serum purified human IgG (IgG) (I4506, Sigma-Aldrich) was prepared in triplicate. Samples were digested with 0.25 μg EndoNF (20:1 sample:EndoNF ratio) in 50 mM sodium phosphate buffer pH 7.5 at 37 °C and 1500 rpm for 2 h. Proteins were then denatured, reduced, alkylated, and precipitated as described.^71, 72^ The precipitated protein pellets were resuspended in 20 μL of 50 mM ammonium bicarbonate and protease digested. rHuNCAM samples were sequentially digested with trypsin (30:1 sample:protease ratio), Asp-N (80:1 sample:protease ratio), and Glu-C (20:1 sample:protease ratio) by incubation in a thermoshaker at 37 °C and 1500 rpm for 16 h. Each protease was deactivated before the addition of the next enzyme by heating at 95 °C for 10 min for trypsin ^73, 74^, and 5 min for Asp-N and Glu-C. IgG samples were digested with trypsin (30:1 sample:protease ratio) by incubation at 37 °C and 1500 rpm for 16 h. Trypsin activity was deactivated by heating at 95 °C for 10 min. Mild acid hydrolysis was performed by acidifying samples with 2% formic acid and heating at 95 °C for 1 h. All samples for MS analysis were desalted with C18 ZipTips (Millipore, USA), dried in a Genevac miVac centrifugal vacuum concentrator, and reconstituted in 0.1% formic acid at a concentration of 50 ng/μL.

### Mass Spectrometry

Samples were analysed with an Orbitrap Elite mass spectrometer (Thermo Fisher Scientific) coupled to an UltiMate 3000 UHPLC system (Thermo Fisher Scientific). 6 μL (300 ng) of sample were injected onto a Vydac Everest C18 column (75 μm × 75 mm, 2 μm, 300 Å, Hichrom, UK) with an Acclaim PepMap C18 trap column (0.3 mm × 5 mm, 5 μm, 100 Å, Thermo Fisher Scientific). The mobile phases were A: 0.1% formic acid, 1% acetonitrile, and B: 0.1 % formic acid, 80% acetonitrile. Loaded peptides were initially washed for 3 min, and then were eluted at a flow rate of 0.3 μL/min in a mobile phase gradient of 3-8% solution B for 5 min, followed by 8-50% solution B for 37 min. Two separate LC-ESI-MS/MS runs were completed with MS1 spectra acquired at *m/z* 300-1800 and *m/z* 700-1800 for rHuNCAM samples or *m/z* 300-1800 and *m/z* 600-1800 for IgG samples. Full width at half-maximum (FWHM) resolution was 120K at *m/z* 400 with an automatic gain control (AGC) target of 1,000,000 and a maximum injection time of 200 ms. The 10 most intense precursors with signal ≥1000 and charge state >1 were selected for MS2 fragmentation using higher-energy collision-induced dissociation (HCD) with an isolation width of 2 Da. MS2 scans were acquired in the Orbitrap mass analyzer with normalized collision energy of 35%, resolution setting of 30K with an AGC target of 100,000, a maximum injection time of 200 ms, and fixed first mass of *m/z* 140. The MS1 *m/z* range for glycopeptides was optimized based on *in silico* digestions and empirical data acquired using MS1 *m/z* 300-1800. The MS1 *m/z* 300-1800 setting was used to calculate occupancy, while the MS1 *m/z* 700-1800 and *m/z* 600-1800 setting was used to calculate the abundance of glycoforms of rHuNCAM and IgG, respectively.

### Data processing

Peptides and glycopeptides were identified using Proteome Discoverer (v2.0.0.802, Thermo Fisher Scientific) with Byonic (v2.13.17, Protein Metrics) as a node to assign peptide-spectrum matches (PSMs). The Byonic search parameters included a maximum of 2 missed cleavages, a precursor mass tolerance of 10 ppm and fragment mass tolerance of 15 ppm. For rHuNCAM, cleavage was set as semi-specific at arginine (R), lysine (K), aspartic acid (D), and glutamic acid (E). For IgG samples cleavage was set as semi-specific at R and K. The peptide output option was set to show all *N*-glycopeptides regardless of score or FDR cutoff as recommended by the manufacturer for simple samples, and decoys were included. Two common modifications and one rare modification were allowed per peptide. Deamidation at asparagine (N) and glutamine (Q), formation of *N*-pyroglutamyl peptide at Q or E, and oxidation of methionine (M) were common modifications; and *N*-glycosylation was a rare modification. Cysteinyl-S-β-propionamide was set as a fixed modification. The glycan database was from Byonic (*N*-glycan 309 mammalian no sodium) with glycans containing NeuGc removed and HexNAc(3)Hex(5)Fuc(1)NeuAc(1) and 16 PSA glycan structures added (Supplementary Table 1). For final calculations, only *N*-glycans with a minimum structure of HexNAc(2)Hex(3) were included. *N*-glycopeptides containing HexNAc or HexNAc(2) were observed, but were not included in further analysis, as assignment of HexNAc or HexNAc(2) can be ambiguous because these structures can exist as *O*-glycans on serine (S) or threonine (T) residues within peptides that contain an *N*-glycosylation sequon. To calculate area under the curve information for precursors the Precursor Ions Area Detector node was run in Proteome Discoverer.

The protein databases contained the amino acid sequence of rHuNCAM (Supplementary Figure 2) or the human proteome database from UniProt (UP000005640 downloaded 20 April 2018; 20,303 reviewed proteins), and a custom contaminants database from Proteome Discoverer and UniProt (304 proteins). PSMs for *N*-glycopeptides were included based on the quality of MS/MS fragmentation with the following minimum requirements: presence of Y1 (peptide + HexNAc) or Y0 (peptide) ions, and presence of relevant HexNAc (*m/z* 204.0866), Hex (*m/z* 163.0601), NeuAc (*m/z* 292.1027) and NeuAc-H_2_O (*m/z* 274.0921) glycan oxonium ions. The relative abundance of each glycoform at each site was calculated as the summed intensity of all *N*-glycopeptides with a particular glycan divided by the summed intensities of all *N*-glycopeptides containing that site. The summed intensity of each glycoform at a site included different peptide proteolysis forms, variable peptide modifications, and all charge states. *N*-glycan occupancy was calculated in samples that had been treated with PNGase F as the summed intensities of deamidated previously glycosylated peptides at a specific site (including all variations of peptide sequences) divided by the summed intensities of all deamidated and unmodified peptides for that site.

Precursor area under the curve data was obtained through the Precursor Ions Area Detector node of Proteome Discoverer. These values were used for PSMs identified through the Byonic search. A python script was used to facilitate high-throughput parsing of unique PSMs and calculation of relative abundances (Supplementary information, https://github.com/bschulzlab/glypnir). The script allowed the user to select the protein of interest, a minimum area under the value, and the number of *N*-glycosylation sites allowed per peptide. The script selected only PSMs with a search engine rank of 1, indicating high confidence PSMs. The unique PSMs selected were sorted based on peptide sequence, modifications, and charge state. Only the highest area under the curve for each unique PSM was chosen for quantification. Each sequon was sorted as either unmodified, deamidated, or glycosylated, and annotated onto the original protein sequence. For glycosylated sequons, the script had an option to group the sequons according to glycan composition. To calculate either occupancy or glycan abundance, the summed area of each unique PSM (based on peptide composition, modifications of the peptide, and peptide charge state) containing the target sequon was divided by the summed area of all unique PSMs containing the target sequon. The output displayed by the script was the occupancy at each sequon or the distribution of glycoforms at each sequon for each sample. All unique glycopeptide PSMs containing sequons of interest used in the computations were manually validated (Supplementary information). The relative abundance of each glycoform at each site was compared between untreated samples and samples treated with EndoNF followed by mild acid hydrolysis using two-tailed, unpaired t-test.

## Results and Discussion

### Generation of polysialylated recombinant HuNCAM

To develop a glycoproteomics workflow capable of measuring site-specific polysialylation, we required a polysialylated glycoprotein substrate. One of the best-studied polysialylated glycoproteins is NCAM,^11, 75^ a single-pass type 1 transmembrane protein with an extracellular domain composed of five Immunoglobulin-like (Ig-like) domains and two fibronectin type III (FN) domains (Figure 2). Human NCAM has six *N*-linked glycosylation sites within the Ig-like domains, and one *N*-linked site close to the transmembrane domain (Figure 2 and Supplementary Figure 2) (https://www.uniprot.org/uniprot/P13591). The 5^th^ and 6^th^ *N*-glycans of native human NCAM are polysialylated (N459 and N488, numbering as per native human protein sequence; N467 and N496 of rHuNCAM) (https://www.uniprot.org/uniprot/P13591). ^11, 22–24^ To obtain polysialylated human NCAM, we expressed a recombinant soluble human NCAM variant (rHuNCAM) in mammalian cells (Figure 2 and Supplementary Figure 2). Polysialylation of rHuNCAM was achieved by co-expression with human B4GALT1, human ST3GAL4, and human ST8SIA4. To verify that the purified rHuNCAM was polysialylated we performed Western blot analysis using anti-PSA and anti-NCAM antibodies with and without pretreatment with EndoNF endosialidase to remove PSA (Figure 3). EndoNF is a tail-spike protein from bacteriophage K1F that cleaves PSA leaving an oligosialic acid chain attached to the glycan.^37^ rHuNCAM showed anti-PSA reactivity (Figure 3, lane 4) that was lost after EndoNF treatment (Figure 3, lane 5), while there was anti-NCAM antibody reactivity in both samples (Figure 3, lanes 6 and 7). We also observed a reduction in the apparent molecular weight of rHuNCAM after EndoNF treatment by Coomassie Blue stained SDS-PAGE (Figure 3, lanes 2 and 3). Together, these results indicated that rHuNCAM was polysialylated. rHuNCAM was subsequently used as our polysialylated glycoprotein substrate for MS workflow development.

**Figure 3.**
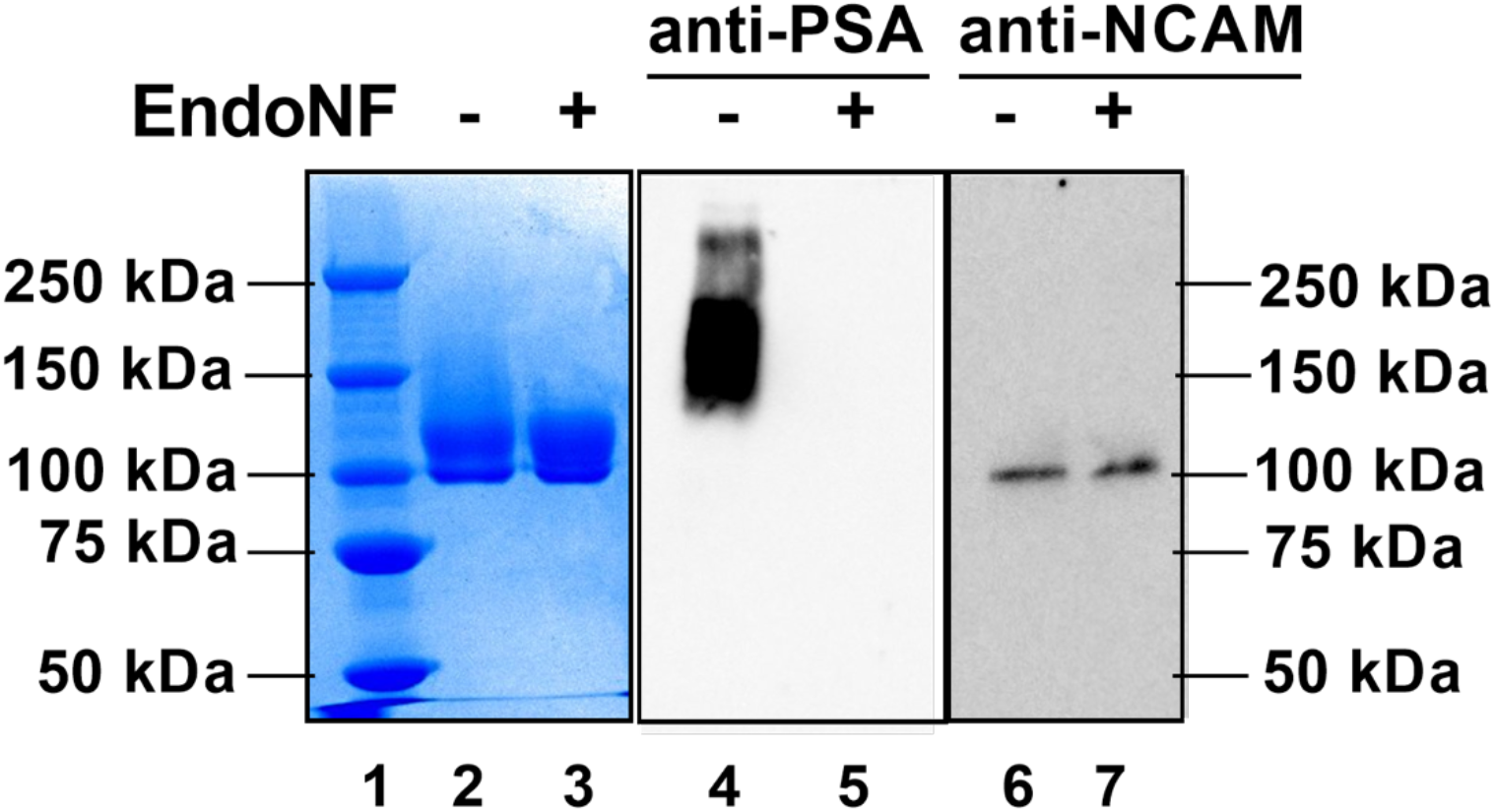
rHuNCAM was partially polysialylated. Purified rHuNCAM was incubated with EndoNF to remove PSA. After incubation, the samples were separated by SDS-PAGE, and either stained with Coomassie Blue (lanes 2 and 3) or transferred to nitrocellulose and used in Western blot analysis (lanes 4-7). PSA and NCAM were detected with anti-PSA (lanes 4-5) and anti-NCAM antibodies (lanes 6-7) respectively.

### An LC-ESI-MS/MS method to measure site-specific polysialylation

Polysialylated glycopeptides are difficult to detect by standard LC-ESI-MS/MS glycoproteomic workflows due to the high molecular weight and negative charge of PSA glycans.^76^ To enable site-specific measurement of polysialylated glycopeptides by LC-ESI MS/MS, PSA must be removed (Figure 1). Indeed, using standard glycoproteomic workflows with only protease digestion and analysis of intact glycopeptides we could not detect PSA-modified glycopeptides from rHuNCAM (Figure 4A and B, Supplementary Tables 2 and 3). We incorporated EndoNF digestion into a standard glycoproteomic workflow as EndoNF can digest PSA to oligosialic acid,^37, 77, 78^ and we had observed that it efficiently removed anti-PSA antibody reactivity from rHuNCAM (Figure 3). We then analysed the types and relative abundance of glycans detected at N467 or N496 in rHuNCAM, which correspond to the 5^th^ and 6^th^ *N*-linked sites of NCAM that are reported to be polysialylated on native NCAM.^11, 22–24^ However, we observed no difference in the relative abundances of glycoforms at these sites in rHuNCAM samples with or without EndoNF digestion (Figure 4A and B, Supplementary Table 2 and 3). The predominant glycoform observed at N467 and N496 in both the untreated and EndoNF treated samples was the oligomannose HexNAc(2)Hex(5) glycan (Figure 4A and B, Supplementary Table 2 and 3). These results suggested that oligosialylated glycopeptides were not detectable by LC-ESI-MS/MS. The negatively charged oligosialic acid chain remaining after EndoNF treatment may reduce the ionisation efficiency of the glycopeptides in positive ion mode, resulting in diminished MS1 signals and limiting detection. To overcome this problem, we treated the samples with mild acid hydrolysis, to completely remove sialic acids.^16, 73, 74^ As expected, after mild acid hydrolysis we could only detect asialylated glycoforms by MS (Figure 4A and B, Supplementary Table 2 and 3). This suggested that EndoNF treatment was not required, and indeed, desialylation with mild acid hydrolysis only resulted in equivalent detection of asialylated glycoforms (Figure 4A and B, Supplementary Table 2 and 3). This treatment also allowed for the detection of several new glycoforms at both N467 and N496 (Figure 4A and B, Supplementary Table 2 and 3). At N467, glycopeptides with HexNAc(3)Hex(4)Fuc(1), HexNAc(3)Hex(5)Fuc(1), HexNAc(3)Hex(6), HexNAc(3)Hex(6)Fuc(1), and HexNAc(4)Hex(5)Fuc(1) glycans were previously undetectable, and increased in abundance after acid hydrolysis (Figure 4A, Supplementary Table 2). Similarly, at the N496 site, glycopeptides with HexNAc(3)Hex(4), HexNAc(3)Hex(5), HexNAc(3)Hex(6), and HexNAc(4)Hex(5) glycans were only detectable after mild acid hydrolysis (Figure 4B and 5, Supplementary Table 3). The new glycoforms detected after hydrolysis (hybrid and complex *N*-glycans) were also the glycan scaffolds that have been previously reported to be polysialylated in bovine NCAM.^64^ Therefore, complete removal of sialic acids by mild acid hydrolysis allowed the detection and relative quantification of polysialylated glycopeptides by LC-ESI-MS/MS.

**Figure 4.**
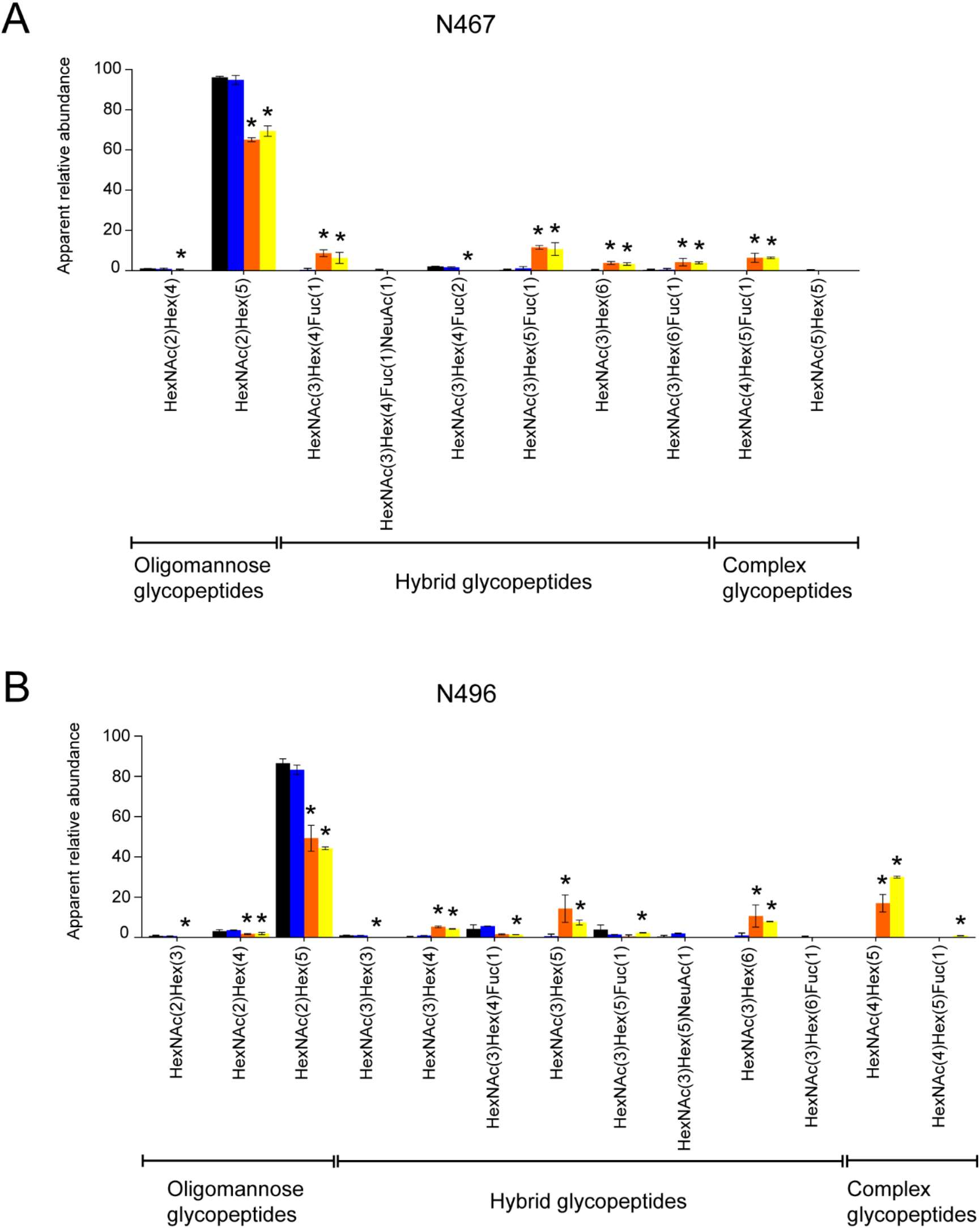
Desialylation allows inference of site specific polysialylation on rHuNCAM. Relative abundance of glycoforms detected by LC-ESI-MS/MS at glycosylation sites **(A)** N467 and **(B)** N496 of rHuNCAM. Black, untreated; blue, EndoNF; orange, EndoNF and mild acid hydrolysis; and yellow, mild acid hydrolysis. Values show mean ± SD, n=3. *, *p* < 0.05 compared to untreated. Full data is shown in Supplementary Tables 2 and 3.

**Figure 5.**
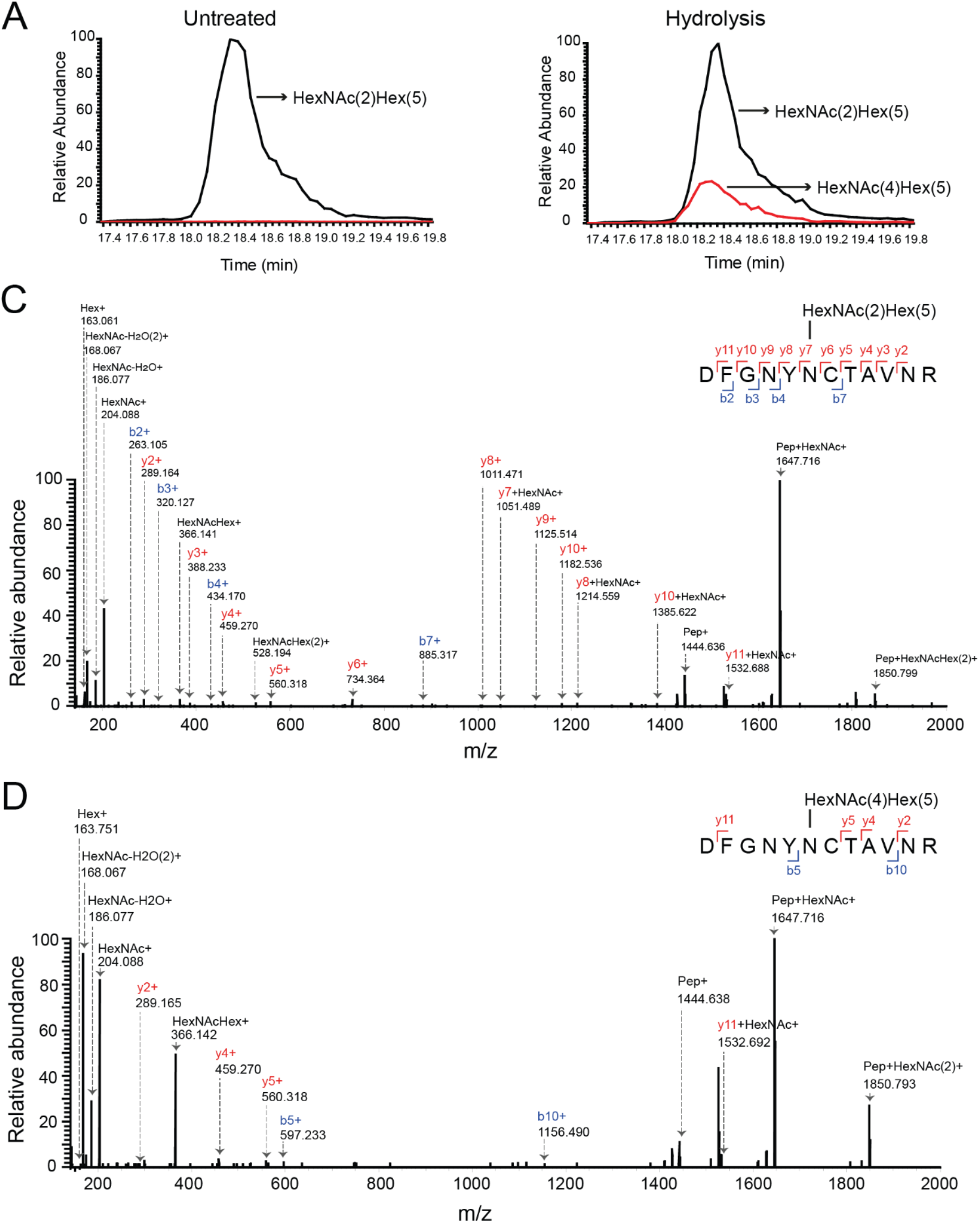
Apparent glycoform abundance of selected DFGNYN^496^CTAVNR glycopeptides of rHuNCAM with and without desialylation. Extracted ion chromatograms of DFGNYNCTAVNR glycopeptide with oligomannose HexNAc(2)Hex(5) (non-polysialylated glycan) or biantennary HexNAc(4)Hex(5) (de-polysialylated glycan) in (**A**) untreated or (**B**) mild acid hydrolysis treated rHuNCAM samples. MS/MS spectra identifying DFGNYNCTAVNR glycopeptides with (**C**) HexNAc(2)Hex(5) or (**D**) HexNAc(4)Hex(5) glycan. All cysteines (C) are alkylated.

To allow focused analysis of glycopeptides containing N467 and N496, the sites known to be polysialylated in native NCAM,^11, 22–24^ we optimized protease digestion and LC-ESI-MS/MS acquisition methods accordingly. However, this generated suboptimal conditions for detection of other *N*-glycopeptides from rHuNCAM. Therefore, to verify that our workflow did not perturb analysis of glycopeptides without PSA, we tested the workflow on serum-purified human immunoglobulin G (IgG), a non-polysialylated glycoprotein (Figure 6, Supplementary Table 4). This analysis focused on IgG2, for which glycopeptides were most robustly detected. IgG2 has one *N*-linked glycosylation site at N176 (https://www.uniprot.org/uniprot/P01859), which is predominantly occupied by bi-antennary glycans.^79^ IgG2 glycoforms observed in the untreated sample included HexNAc(4)Hex(3)Fuc(1), HexNAc(4)Hex(4)Fuc(1), HexNAc(4)Hex(5)Fuc(1), HexNAc(4)Hex(4)Fuc(1)NeuAc(1), and HexNAc(4)Hex(5)Fuc(1)NeuAc(1) (Figure 6, Supplementary Table 4). This glycoform distribution is similar to previous reports.^79^ After EndoNF treatment, we did not detect any new glycoforms on IgG2, and there was no significant change in the relative abundance of the observed glycoforms compared to the untreated samples (Figure 6, Supplementary Table 4). This result was expected, given that IgG2 is not reported to contain PSA-modified glycans. Importantly, the only changes observed in IgG2 glycoform structure and abundance in samples treated with EndoNF and mild acid hydrolysis compared to untreated samples were the disappearance of previously monosialylated structures, and the increase in abundance of the corresponding unsialylated forms of these structures (Figure 6, Supplementary Table 4). For example, we observed the disappearance of the HexNAc(4)Hex(4)Fuc(1)NeuAc(1) and HexNAc(4)Hex(5)Fuc(1)NeuAc(1) glycoforms in desialylated samples, and an increase in the abundance of the corresponding HexNAc(4)Hex(4)Fuc(1) and HexNAc(4)Hex(5)Fuc(1) structures (Figure 6, Supplementary Table 4). We did not detect any new glycoform in the desialylated sample. Since the relative abundance of nonsialylated glycoforms increases after desialylation, our workflow is more effective when comparing potentially polysialylated glycoproteins with their non-polysialylated counterparts. We also note that some defucosylation was observed with mild acid hydrolysis (Figure 4 and Supplementary Tables 2 and 3), although the extent of fucose loss did not substantially impact determination of the site-specificity and occupancy of polysialylation. Together, these results show that our workflow is a useful method to detect site-specific polysialylation in glycoproteins.

**Figure 6.**
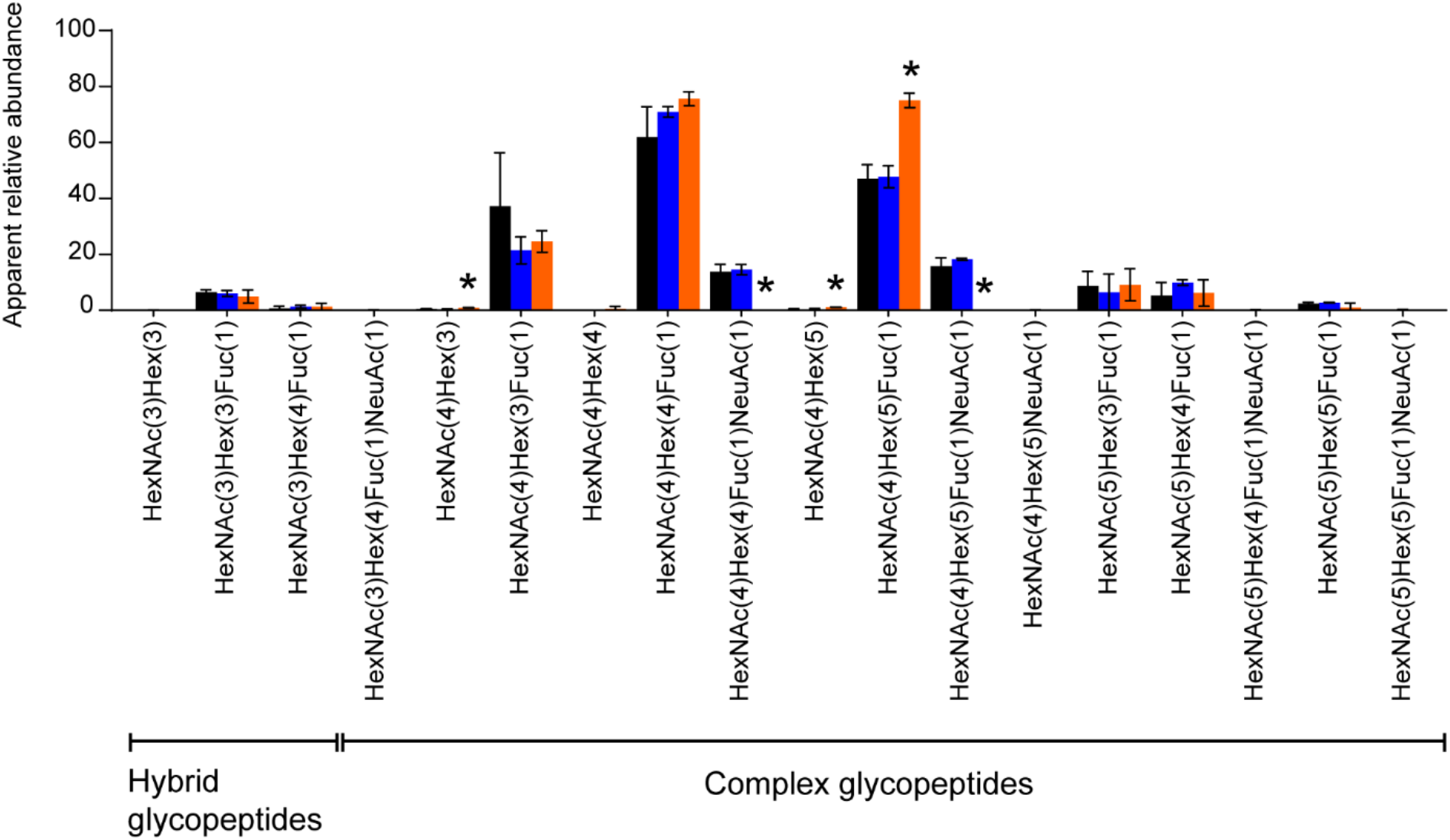
Desialylation does not strongly affect the glycan structures on human serum IgG. Relative abundance of glycoforms detected by LC-ESI-MS/MS at the glycosylation site N176 of IgG2. Black, untreated; blue, EndoNF; orange, EndoNF and mild acid hydrolysis. Values show mean ± SD, n=3. *, *p* < 0.05 compared to untreated. Full data is shown in Supplementary Table 4.

## Conclusions

PSA has key physiological roles and therapeutic applications. To facilitate the study of polysialylation and the development of new PSA technologies, we have developed an LC-ESI-MS/MS method that allows for the detection and measurement of site-specific polysialylation. Our method combines carefully selected proteases to generate peptides of appropriate length for LC-ESI-MS/MS detection with mild acid hydrolysis to eliminate sialic acids and newly developed Python scripts for data processing and calculation of glycoform abundance. We have demonstrated that this method distinguishes PSA- and non-PSA modified glycopeptides, can identify the precise peptides modified by PSA, and allows measurement of the site-specific extent of polysialylation.

## Acknowledgements

We thank Dr Amanda Nouwens and Peter Josh at The University of Queensland, School of Chemistry and Molecular Biosciences Mass Spectrometry Facility for their assistance and expertise.

## References

1. Colley, K. J.; Varki, A.; Kinoshita, T., Cellular Organization of Glycosylation. In Essentials of Glycobiology, rd; Varki, A.; Cummings, R. D.; Esko, J. D.; Stanley, P.; Hart, G. W.; Aebi, M.; Darvill, A. G.; Kinoshita, T.; Packer, N. H.; Prestegard, J. H.; Schnaar, R. L.; Seeberger, P. H., Eds. Cold Spring Harbor (NY), 2015; pp 41–49.

2. Zacchi, L. F.; Caramelo, J. J.; McCracken, A. A.; Brodsky, J. L., Endoplasmic Reticulum-Associated Degradation and Protein Quality Control. In Encyclopedia of Cell Biology, Stahl, R. A. B. a. P. D., Ed. Academic Press: 2016; Vol. 1, pp 596–611.

3. Helenius, A.; Aebi, M., Roles of N-linked glycans in the endoplasmic reticulum. Annu Rev Biochem 2004, 73, 1019–1049.

4. Caramelo, J. J.; Parodi, A. J., A sweet code for glycoprotein folding. FEBS Lett 2015, 589 (22), 3379–87.

5. Stanley, P.; Taniguchi, N.; Aebi, M., N-Glycans. In Essentials of Glycobiology, rd; Varki, A.; Cummings, R. D.; Esko, J. D.; Stanley, P.; Hart, G. W.; Aebi, M.; Darvill, A. G.; Kinoshita, T.; Packer, N. H.; Prestegard, J. H.; Schnaar, R. L.; Seeberger, P. H., Eds. Cold Spring Harbor (NY), 2015; pp 99–111.

6. Rini, J.; Esko, J.; Varki, A., Chapter 5 Glycosyltransferases and Glycan-processing Enzymes. In Essentials of Glycobiology [Internet], 3rd ed.; A, V.; Cummings, R.; Jd, E.; al., e., Eds. Cold Spring Harbor Laboratory Press: Cold Spring Harbor (NY), 2015–2017.

7. Rudd, P. M.; Dwek, R. A., Glycosylation: heterogeneity and the 3D structure of proteins. Crit Rev Biochem Mol Biol 1997, 32 (1), 1–100.

8. Zacchi, L. F.; Schulz, B. L., N-glycoprotein macroheterogeneity: biological implications and proteomic characterization. Glycoconj J 2016, 33 (3), 359–76.

9. Varki, A.; Schnaar, R. L.; Schauer, R., Sialic Acids and Other Nonulosonic Acids. In Essentials of Glycobiology, rd; Varki, A.; Cummings, R. D.; Esko, J. D.; Stanley, P.; Hart, G. W.; Aebi, M.; Darvill, A. G.; Kinoshita, T.; Packer, N. H.; Prestegard, J. H.; Schnaar, R. L.; Seeberger, P. H., Eds. Cold Spring Harbor (NY), 2015; pp 179–195.

10. Sato, C.; Kitajima, K., Disialic, oligosialic and polysialic acids: distribution, functions and related disease. The Journal of Biochemistry 2013, 154 (2), 115–136.

11. Colley, K. J.; Kitajima, K.; Sato, C., Polysialic acid: biosynthesis, novel functions and applications. Crit Rev Biochem Mol Biol 2014, 49 (6), 498–532.

12. Finne, J.; Finne, U.; Deagostini-Bazin, H.; Goridis, C., Occurrence of alpha 2-8 linked polysialosyl units in a neural cell adhesion molecule. Biochemical and biophysical research communications 1983, 112 (2), 482–7.

13. Sebastian, P. G.; Manuela, R.; Moritz, K.; Katinka, E.; Imke, O.-N.; Miriam, S.; Maike, H.; Birgit, W.; Herbert, H.; Rudolf, G.; Martina, M.; Hildegard, G., Synaptic cell adhesion molecule SynCAM 1 is a target for polysialylation in postnatal mouse brain. Proceedings of the National Academy of Sciences 2010, 107 (22), 10250.

14. Curreli, S.; Arany, Z.; Gerardy-Schahn, R.; Mann, D.; Stamatos, N. M., Polysialylated Neuropilin-2 Is Expressed on the Surface of Human Dendritic Cells and Modulates Dendritic Cell-T Lymphocyte Interactions. Polysialylated Neuropilin-2 Is Expressed on the Surface of Human Dendritic Cells and Modulates Dendritic Cell-T Lymphocyte Interactions 2007, 282 (42), 30346–30356.

15. Zuber, C.; Lackie, P. M.; Catterall, W. A.; Roth, J., Polysialic acid is associated with sodium channels and the neural cell adhesion molecule N-CAM in adult rat brain. The Journal of biological chemistry 1992, 267 (14), 9965–9971.

16. James, W. M.; Agnew, W. S., Multiple oligosaccharide chains in the voltage-sensitive Na channel from electrophorus electricus: evidence for alpha-2,8-linked polysialic acid. Biochem Biophys Res Commun 1987, 148 (2), 817–26.

17. Kiermaier, E.; Moussion, C.; Veldkamp, C. T.; Gerardy-Schahn, R.; de Vries, I.; Williams, L. G.; Chaffee, G. R.; Phillips, A. J.; Freiberger, F.; Imre, R.; Taleski, D.; Payne, R. J.; Braun, A.; Forster, R.; Mechtler, K.; Muhlenhoff, M.; Volkman, B. F.; Sixt, M., Polysialylation controls dendritic cell trafficking by regulating chemokine recognition. Science 2016, 351 (6269), 186–90.

18. Werneburg, S.; Buettner, F. F.; Erben, L.; Mathews, M.; Neumann, H.; Muhlenhoff, M.; Hildebrandt, H., Polysialylation and lipopolysaccharide-induced shedding of E-selectin ligand-1 and neuropilin-2 by microglia and THP-1 macrophages. Glia 2016, 64 (8), 1314–30.

19. Yabe, U.; Sato, C.; Matsuda, T.; Kitajima, K., Polysialic acid in human milk. CD36 is a new member of mammalian polysialic acid-containing glycoprotein. The Journal of biological chemistry 2003, 278 (16), 13875.

20. Muhlenhoff, M.; Eckhardt, M.; Bethe, A.; Frosch, M.; Gerardy-Schahn, R., Autocatalytic polysialylation of polysialyltransferase-1. The EMBO journal 1996, 15 (24), 6943–50.

21. Close, B. E.; Colley, K. J., In vivo autopolysialylation and localization of the polysialyltransferases PST and STX. The Journal of biological chemistry 1998, 273 (51), 34586–93.

22. Nelson, R. W.; Bates, P. A.; Rutishauser, U., Protein determinants for specific polysialylation of the neural cell adhesion molecule. The Journal of biological chemistry 1995, 270 (29), 17171.

23. Close, B. E.; Mendiratta, S. S.; Geiger, K. M.; Broom, L. J.; Ho, L.-L.; Colley, K. J., The minimal structural domains required for neural cell adhesion molecule polysialylation by PST/ST8Sia IV and STX/ST8Sia II. The Journal of biological chemistry 2003, 278 (33), 30796.

24. Foley, D. A.; Swartzentruber, K. G.; Lavie, A.; Colley, K. J., Structure and Mutagenesis of Neural Cell Adhesion Molecule Domains Evidence for Flexibility in the Placement of Polysialic Acid Attachment Sites. J. Biol. Chem. 2010, 285 (35).

25. Sato, C.; Kitajima, K., Impact of structural aberrancy of polysialic acid and its synthetic enzyme ST8SIA2 in schizophrenia. Front. Cell. Neurosci. 2013, 7.

26. Barbeau, D.; Liang, J. J.; Robitaille, Y.; Quirion, R.; Srivastava, L. K., Decreased Expression of the Embryonic Form of the Neural Cell Adhesion Molecule in Schizophrenic Brains. Proceedings of the National Academy of Sciences of the United States of America 1995, 92 (7), 2785–2789.

27. Sara, M. E.; Simon, J. A.; Maria, S.; Haneen, A. B.; Goreti Ribeiro, M.; Paul, M. L.; Klaus, P.; Robert, A. F., Polysialic acid sustains cancer cell survival and migratory capacity in a hypoxic environment. Scientific Reports 2016, 6 (1).

28. El Maarouf, A.; Rutishauser, U., Use of PSA-NCAM in repair of the central nervous system. Advances in Experimental Medicine and Biology 2010, 663, 137–147.

29. Bader, R. A.; Wardwell, P. R., Polysialic acid: overcoming the hurdles of drug delivery. Therapeutic delivery 2014, 5 (3), 235–7.

30. Wang, X.-J.; Gao, Y.-P.; Lu, N.-N.; Li, W.-S.; Xu, J.-F.; Ying, X.-Y.; Wu, G.; Liao, M.-H.; Tan, C.; Shao, L.-X.; Lu, Y.-M.; Zhang, C.; Fukunaga, K.; Han, F.; Du, Y.-Z., Endogenous Polysialic Acid Based Micelles for Calmodulin Antagonist Delivery against Vascular Dementia. ACS applied materials & interfaces 2016, 8 (51), 35045–35058.

31. Zhang, R.; Jain, S.; Rowland, M.; Hussain, N.; Agarwal, M.; Gregoriadis, G., Development and testing of solid dose formulations containing polysialic acid insulin conjugate: next generation of long-acting insulin. Journal of diabetes science and technology 2010, 4 (3), 532.

32. Kontermann, R. E., Strategies for extended serum half-life of protein therapeutics. Current Opinion in Biotechnology 2011, 22 (6), 868–876.

33. Fernandes, A. I.; Gregoriadis, G., The effect of polysialylation on the immunogenicity and antigenicity of asparaginase: implication in its pharmacokinetics. International Journal of Pharmaceutics 2001, 217 (1), 215–224.

34. Lindhout, T.; Iqbal, U.; Willis, L. M.; Reid, A. N.; Li, J.; Liu, X.; Moreno, M.; Wakarchuk, W. W., Site-specific enzymatic polysialylation of therapeutic proteins using bacterial enzymes. Proc Natl Acad Sci U S A 2011, 108 (18), 7397–402.

35. Frosch, M.; Görgen, I.; Boulnois, G. J.; Timmis, K. N.; Bitter-Suermann, D., NZB mouse system for production of monoclonal antibodies to weak bacterial antigens: isolation of an IgG antibody to the polysaccharide capsules of Escherichia coli K1 and group B meningococci. Proceedings of the National Academy of Sciences of the United States of America 1985, 82 (4), 1194.

36. Sato, C.; Kitajima, K.; Inoue, S.; Seki, T.; Troy Ii, F. A.; Inoue, Y., Characterization of the antigenic specificity of four different anti-(α2→8-linked polysialic acid) antibodies using lipid-conjugated oligo/polysialic acids. Journal of Biological Chemistry 1995, 270 (32), 18923–18928.

37. Morley, T. J.; Willis, L. M.; Whitfield, C.; Wakarchuk, W. W.; Withers, S. G., New Sialidase Mechanism: BACTERIOPHAGE K1F ENDO-SIALIDASE IS AN INVERTING GLYCOSIDASE. New Sialidase Mechanism: BACTERIOPHAGE K1F ENDO-SIALIDASE IS AN INVERTING GLYCOSIDASE 2009, 284 (26), 17404–17410.

38. Katharina, S.; Achim, D.; Martina, M.; Rita, G.-S.; Ralf, F., Crystal structure of the polysialic acid–degrading endosialidase of bacteriophage K1F. Nature Structural & Molecular Biology 2004, 12 (1), 90.

39. Aalto, J.; Pelkonen, S.; Kalimo, H.; Finne, J., Mutant bacteriophage with non-catalytic endosialidase binds to both bacterial and eukaryotic polysialic acid and can be used as probe for its detection. Official Journal of the International Glycoconjugate Organization 2001, 18 (10), 751–758.

40. Kitajima, K.; Varki, N.; Sato, C., Advanced Technologies in Sialic Acid and Sialoglycoconjugate Analysis. Top Curr Chem 2015, 367, 75–103.

41. Galuska, S. P., Chapter 13: Advances in Sialic Acid and Polysialic Acid Detection Methodologies. In Sialobiology: Structure, Biosynthesis and Function, Martinez-Duncker, J. T. a. I., Ed. Bentham Science Publishers: 2013; pp 448–475.

42. Inoue, S.; Inoue, Y., A challenge to the ultrasensitive chemical method for the analysis of oligo- and polysialic acids at a nanogram level of colominic acid and a milligram level of brain tissues. Biochimie 2001, 83 (7), 605–613.

43. Inoue; Lin, S. L.; Lee, Y. C.; Inoue, An ultrasensitive chemical method for polysialic acid analysis. Glycobiology 2001, 11 (9), 759–767.

44. Inoue, S.; Inoue, Y., Ultrasensitive analysis of sialic acids and oligo/polysialic acids by fluorometric high-performance liquid chromatography. Methods in enzymology 2003, 362, 543–60.

45. Sato, C.; Inoue, S.; Matsuda, T.; Kitajima, K., Fluorescent-Assisted Detection of Oligosialyl Units in Glycoconjugates. Analytical Biochemistry 1999, 266 (1), 102–109.

46. Sato, C.; Inoue, S.; Matsuda, T.; Kitajima, K., Development of a highly sensitive chemical method for detecting alpha2⤍8-linked oligo/polysialic acid residues in glycoproteins blotted on the membrane. Anal Biochem 1998, 261 (2), 191–7.

47. Rohr, T. E.; Troy, F. A., Structure and biosynthesis of surface polymers containing polysialic acid in Escherichia coli. J Biol Chem 1980, 255 (6), 2332–42.

48. Zhang, Y.; Inoue, Y.; Inoue, S.; Lee, Y. C., Separation of Oligo/Polymers of 5-N-Acetylneuraminic Acid, 5-N-Glycolylneuraminic Acid, and 2-Keto-3-deoxy-d-glycero-d-galacto-nononic Acid by High-Performance Anion-Exchange Chromatography with Pulsed Amperometric Detector. Analytical Biochemistry 1997, 250 (2), 245–251.

49. Shu-Ling, L.; Inoue, Y.; Inoue, S., Evaluation of high-performance anion-exchange chromatography with pulsed electrochemical and fluorometric detection for extensive application to the analysisof homologous series of oligo- and polysialic acids in bioactive molecules. Glycobiology 1999, 9 (8), 807.

50. Michon, F.; Brisson, J. R.; Jennings, H. J., Conformational Differences between Linear α(2→8)-Linked Homosialooligosaccharides and the Epitope of the Group B Meningococcal Polysaccharide. Biochemistry 1987, 26 (25), 8399–8405.

51. Inoue, S.; Iwasaki, M., Isolation of a novel glycoprotein from the eggs of rainbow trout: Occurrence of disialosyl groups on all carbohydrate chains. Biochemical and Biophysical Research Communications 1978, 83 (3), 1018–1023.

52. Ijuin, T.; Kitajima, K.; Song, Y.; Kitazume, S.; Inoue, S.; Haslam, S.; Morris, H.; Dell, A.; Inoue, Y., Isolation and identification of novel sulfated and nonsulfated oligosialyl glycosphingolipids from sea urchin sperm. Official Journal of the International Glycoconjugate Organization 1996, 13 (3), 401–413.

53. Kitazume, S.; Kitajima, K.; Inoue, S.; Troy, F. A., 2nd; Cho, J. W.; Lennarz, W. J.; Inoue, Y., Identification of polysialic acid-containing glycoprotein in the jelly coat of sea urchin eggs. Occurrence of a novel type of polysialic acid structure. The Journal of biological chemistry 1994, 269 (36), 22712–8.

54. Thaysen-Andersen, M.; Larsen, M. R.; Packer, N. H.; Palmisano, G., Structural analysis of glycoprotein sialylation – Part I: pre-LC-MS analytical strategies RSC Advances 2013, 3 (45), 22683–22705.

55. Palmisano, G.; Larsen, M. R.; Packer, N. H.; Thaysen-Andersen, M., Structural analysis of glycoprotein sialylation – part II: LC-MS based detection. RSC Advances 2013, 3 (45), 22706–22726.

56. Galuska, C. E.; Maass, K.; Galuska, S. P., Mass Spectrometric Analysis of Oligo- and Polysialic Acids. Methods in molecular biology (Clifton, N.J.) 2015, 1321, 417–26.

57. Galuska, S. P.; Geyer, H.; Mink, W.; Kaese, P.; Kuhnhardt, S.; Schafer, B.; Muhlenhoff, M.; Freiberger, F.; Gerardy-Schahn, R.; Geyer, R., Glycomic strategy for efficient linkage analysis of di-, oligo- and polysialic acids. Journal of proteomics 2012, 75 (17), 5266–78.

58. Galuska, S.; Geyer, R.; Muhlenhoff, M.; Geyer, H., Characterization of oligo- and polysialic acids by MALDI-TOF-MIS. Anal. Chem. 2007, 79 (18), 7161–7169.

59. Galuska, S. P.; Geyer, H.; Bleckmann, C.; Rohrich, R. C.; Maass, K.; Bergfeld, A. K.; Muhlenhoff, M.; Geyer, R., Mass spectrometric fragmentation analysis of oligosialic and polysialic Acids.(Author abstract). Analytical Chemistry 2010, 82 (5), 2059.

60. Miyata, S.; Sato, C.; Kitamura, S.; Toriyama, M.; Kitajima, K., A major flagellum sialoglycoprotein in sea urchin sperm contains a novel polysialic acid, an 2,9-linked poly-N-acetylneuraminic acid chain, capped by an 8-O -sulfated sialic acid residue. Glycobiology 2004, 14 (9), 827–840.

61. Kronewitter, S. R.; Marginean, I.; Cox, J. T.; Zhao, R.; Hagler, C. D.; Shukla, A. K.; Carlson, T. S.; Adkins, J. N.; Camp, D. G., 2nd; Moore, R. J.; Rodland, K. D.; Smith, R. D., Polysialylated N-glycans identified in human serum through combined developments in sample preparation, separations, and electrospray ionization-mass spectrometry. Anal Chem 2014, 86 (17), 8700–10.

62. Rollenhagen, M.; Buettner, F. F.; Reismann, M.; Jirmo, A. C.; Grove, M.; Behrens, G. M.; Gerardy-Schahn, R.; Hanisch, F. G.; Muhlenhoff, M., Polysialic acid on neuropilin-2 is exclusively synthesized by the polysialyltransferase ST8SiaIV and attached to mucin-type o-glycans located between the b2 and c domain. J Biol Chem 2013, 288 (32), 22880–92.

63. Liedtke, S.; Geyer, H.; Wuhrer, M.; Geyer, R.; Frank, G.; Gerardy-Schahn, R.; Zahringer, U.; Schachner, M., Characterization of N-glycans from mouse brain neural cell adhesion molecule. Glycobiology 2001, 11 (5), 373–384.

64. Von Der Ohe, M.; Wheeler, S.; Wuhrer, M.; Harvey, D.; Liedtke, S.; Muhlenhoff, M.; Gerardy-Schahn, R.; Geyer, H.; Dwek, R.; Geyer, R.; Wing, D.; Schachner, M., Localization and characterization of polysialic acid-containing N-linked glycans from bovine NCAM. Glycobiology 2002, 12 (1), 47–47.

65. Chen, C.-G.; Fabri, L. J.; Wilson, M. J.; Panousis, C., One-step zero-background IgG reformatting of phage-displayed antibody fragments enabling rapid and high-throughput lead identification. Nucleic acids research 2014, 42 (4), e26–e26.

66. Pham, P. L.; Perret, S.; Cass, B.; Carpentier, E.; St-Laurent, G.; Bisson, L.; Kamen, A.; Durocher, Y., Transient gene expression in HEK293 cells: Peptone addition posttransfection improves recombinant protein synthesis. Biotechnology and Bioengineering 2005, 90 (3), 332–344.

67. Schmidt, P. M.; Abdo, M.; Butcher, R. E.; Yap, M.-Y.; Scotney, P. D.; Ramunno, M. L.; Martin-Roussety, G.; Owczarek, C.; Hardy, M. P.; Chen, C.-G.; Fabri, L. J., A robust robotic high-throughput antibody purification platform. Journal of Chromatography A 2016, 1455, 9–19.

68. Panousis, C.; Dhagat, U.; Edwards, K. M.; Rayzman, V.; Hardy, M. P.; Braley, H.; Gauvreau, G. M.; Hercus, T. R.; Smith, S.; Sehmi, R.; McMillan, L.; Dottore, M.; McClure, B. J.; Fabri, L. J.; Vairo, G.; Lopez, A. F.; Parker, M. W.; Nash, A. D.; Wilson, N. J.; Wilson, M. J.; Owczarek, C. M., CSL311, a novel, potent, therapeutic monoclonal antibody for the treatment of diseases mediated by the common β chain of the IL-3, GM-CSF and IL-5 receptors. mAbs 2016, 8 (3).

69. Schaer, C. A.; Owczarek, C.; Deuel, J. W.; Schauer, S.; Baek, J. H.; Yalamanoglu, A.; Hardy, M. P.; Scotney, P. D.; Schmidt, P. M.; Pelzing, M.; Soupourmas, P.; Buehler, P. W.; Schaer, D. J., Phenotype-specific recombinant haptoglobin polymers co-expressed with C1r-like protein as optimized hemoglobin-binding therapeutics. BMC Biotechnology 2018, 18 (1), <xocs:firstpage xmlns:xocs=“”/>.

70. Jamaluddin, M. F. B.; Bailey, U.-M.; Tan, N. Y.; Stark, A. P.; Schulz, B. L., Polypeptide binding specificities of Saccharomyces cerevisiae oligosaccharyltransferase accessory proteins Ost3p and Ost6p. Protein Science 2011, 20 (5), 849–855.

71. Nguyen, L. T.; Zacchi, L. F.; Schulz, B. L.; Moore, S. S.; Fortes, M. R. S., Adipose tissue proteomic analyses to study puberty in Brahman heifers. Journal of Animal Science 2018, 96 (6), 2392–2398.

72. Bailey, U.-M.; Punyadeera, C.; Cooper-White, J. J.; Schulz, B. L., Analysis of the extreme diversity of salivary alpha-amylase isoforms generated by physiological proteolysis using liquid chromatography–tandem mass spectrometry. Journal of Chromatography B 2012, 911, 21–26.

73. Varki, A.; Diaz, S., The release and purification of sialic acids from glycoconjugates: Methods to minimize the loss and migration of O-acetyl groups. Analytical Biochemistry 1984, 137 (1), 236–247.

74. Manzi, A., Acid hydrolysis for release of monosaccharides. Current protocols in molecular biology / edited by Frederick M. Ausubel … [et al.] 2001, 17, Unit17.16.

75. Berezin, V., Structure and Function of the Neural Cell Adhesion Molecule NCAM. New York, NY: Springer New York: 2010.

76. Nwosu, C. C.; Strum, J. S.; An, H. J.; Lebrilla, C. B., Enhanced detection and identification of glycopeptides in negative ion mode mass spectrometry. Anal Chem 2010, 82 (23), 9654–62.

77. Jakobsson, E.; Schwarzer, D.; Jokilammi, A.; Finne, J., Endosialidases: Versatile Tools for the Study of Polysialic Acid. Top Curr Chem 2015, 367, 29–73.

78. Schulz, E. C.; Schwarzer, D.; Frank, M.; Stummeyer, K.; Mühlenhoff, M.; Dickmanns, A.; Gerardy-Schahn, R.; Ficner, R., Structural Basis for the Recognition and Cleavage of Polysialic Acid by the Bacteriophage K1F Tailspike Protein EndoNF. Journal of Molecular Biology 2010, 397 (1), 341–351.

79. Wuhrer, M.; Stam, J. C.; van de Geijn, F. E.; Koeleman, C. A.; Verrips, C. T.; Dolhain, R. J.; Hokke, C. H.; Deelder, A. M., Glycosylation profiling of immunoglobulin G (IgG) subclasses from human serum. Proteomics 2007, 7 (22), 4070–81.

80. Gerardy-Schahn, R.; Delannoy, P.; von Itzstein, M., SialoGlyco Chemistry and Biology II Tools and Techniques to Identify and Capture Sialoglycans. 1st ed. 2015.․ ed.; Cham : Springer International Publishing : Imprint: Springer: 2015.

81. Schwarzer, D.; Stummeyer, K.; Freiberger, F.; Grove, M.; Mühlenhoff, M.; Haselhorst, T.; Von Itzstein, M.; Gerady-Schahn, R.; Rode, B.; Scheper, T., Proteolytic release of the intramolecular chaperone domain confers processivity to endosialidase F. Journal of Biological Chemistry 2009, 284 (14), 9465–9474.

82. Finne, J., Occurrence of unique polysialosyl carbohydrate units in glycoproteins of developing brain. Journal of Biological Chemistry 1982, 257 (20), 11966–11970.

