## Supplementary Information for "Glycoproteomic measurement of site-specific polysialylation"

### Supplementary Figures

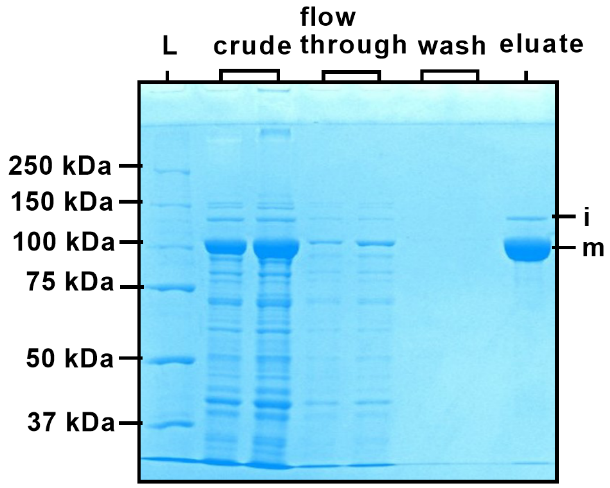


**Supplementary Figure S1. Expression and purification of EndoNF.** EndoNF expressed in an *E. coli* Rosetta strain was purified using amylose-agarose resin. Aliquots from the crude, flow through, wash, and eluate samples were separated by SDS-PAGE and the gel was stained with Coomassie Blue. EndoNF appeared as two bands in the eluate. The most prominent band was ~ 118 kDa (marked 'm'), corresponding to the mature EndoNF enzyme; while the top band of ~ 136 kDa corresponded to the immature EndoNF (marked 'i') that still contains the C-terminal chaperone domain (verified by in gel digestion and mass spectrometry analysis (data not shown)) which becomes self-cleaved during maturation [80, 81].

MLQTKDLIWTLFFLGTAVSDYKDDDDKLQVDIVPSQGEISVGESKFFLCQVAGDA<br>
KDKDISWFSPNGEKLTPNQQRISVVWNDDSSSTLTIYNANIDDAGIYKCVVTGED<br>
GSESEATVNVKIFQKLMFKNAPTPQEFREGEDAVIVCDVVSSLPPTIIWKHKGRD<br>
VILKKDVRFIVLSNNYLQIRGIKKTDEGTYRCEGRILARGEINFKDIQVIVNVPP<br>
TIQARQNIV**N**$^{230}$ATANLGQSVTLVCDAEGFPEPTMSWTKDGEQIEQEEDDEKYIF<br>
SDDSSQLTIKKVDKNDEAEYICIAENKAGEQDATIHLKVFAKPKITYVE**N**$^{323}$QTA<br>
MELEEQVTLTCEASGDPIPSITWRTSTR**N**$^{355}$ISSEEKASWTRPEKQETLDGHMVV<br>
RSHARVSSLTLKSIQYTDAGEYICTASNTIGQDSQSMYLEVQYAPKLQGPVAVYT<br>
WEGNQV**N**$^{441}$ITCEVFAYPSATISWFRDGQLLPSS**N**$^{467}$YSNIKIYNTPSASYLEVT<br>
PDSENDFGNY**N**$^{496}$CTAVNRIGQESLEFILVQADTPSSPSIDQVEPYSSTAQVQFD<br>
EPEATGGVPILKYKAEWRAVGEEVWHSKWYDAKEASMEGIVTIVGLKPETTYAVR<br>
LAALNGKGLGEISAASEFKTQPVQGEPSAPKLEGQMGEDGNSIKVNLIKQDDGGS<br>
PIRHYLVRYRALSSEWKPEIRLPSGSDHVMLKSLDWNAEYEVYVVAENQQGKSKA<br>
AHFVFRTSAQPTAIPA**N**$^{720}$GSPTSGHHHHHHHH

**Supplementary Figure S2. Amino acid sequence of the HuNCAM(1-19)-FLAG-HuNCAM(20-718)-8His used in this study**. The amino acid sequence of rHuNCAM consists of an endogenous signal peptide (green), followed by a FLAG tag (purple), then the NCAM open reading frame truncated at amino acid 718 to remove the transmembrane domain, and finally an 8His tag attached to the C-terminal end of rHuNCAM (yellow). The asparagine residues within the *N*-linked glycosylation sites are highlighted in blue (no polysialylation) or red (polysialylated) based on previous reports [22, 23, 82]

### Supplementary Tables

| **Supplementary Table S1. Additional polysialylated *N*-glycan structures added to the Byonic database** |
|---|
| **HexNAc(4)Hex(5)Fuc(1)NeuAc(3)** |
| **HexNAc(4)Hex(5)Fuc(1)NeuAc(4)** |
| **HexNAc(4)Hex(5)Fuc(1)NeuAc(5)** |
| **HexNAc(4)Hex(5)Fuc(1)NeuAc(6)** |
| **HexNAc(4)Hex(5)Fuc(1)NeuAc(7)** |
| **HexNAc(4)Hex(5)Fuc(1)NeuAc(8)** |
| **HexNAc(4)Hex(5)Fuc(1)NeuAc(9)** |
| **HexNAc(4)Hex(5)Fuc(1)NeuAc(10)** |
| **HexNAc(4)Hex(5)NeuAc(3)** |
| **HexNAc(4)Hex(5)NeuAc(4)** |
| **HexNAc(4)Hex(5)NeuAc(5)** |
| **HexNAc(4)Hex(5)NeuAc(6)** |
| **HexNAc(4)Hex(5)NeuAc(7)** |
| **HexNAc(4)Hex(5)NeuAc(8)** |
| **HexNAc(4)Hex(5)NeuAc(9)** |
| **HexNAc(4)Hex(5)NeuAc(10)** |

**Supplementary Table S2. Glycoform abundance at N467 in rHuNCAM in untreated, EndoNF treated, EndoNF and mild acid hydrolysis treated, and mild acid hydrolysis treated samples. *, $p < 0.05$ compared to untreated.**

| | **Untreated** | **EndoNF** | **EndoNF and mild acid hydrolysis** | **Mild acid Hydrolysis** |
|---|---|---|---|---|
| **Glycoforms** | **Mean ± SD** | **Mean ± SD** | **Mean ± SD** | **Mean ± SD** |
| **HexNAc(2)Hex(4)** | 1.1 ± 0.1 | 0.9 ± 0.4 | 0.2 ± 0.4* | * |
| **HexNAc(2)Hex(5)** | 96.1 ± 0.5 | 94.8 ± 2.2 | 65.1 ± 1.0* | 69.4 ± 2.6* |
| **HexNAc(3)Hex(4)Fuc(1)** | | 0.4 ± 0.7 | 8.7 ± 1.7* | 6.3 ± 2.7* |
| **HexNAc(3)Hex(4)Fuc(1)NeuAc(1)** | | 0.2 ± 0.4 | | |
| **HexNAc(3)Hex(4)Fuc(2)** | 2.1 ± 0.2 | 1.6 ± 0.3 | * | * |
| **HexNAc(3)Hex(5)Fuc(1)** | 0.4 ± 0.3 | 1.2 ± 0.9 | 11.6 ± 1.0* | 10.7 ± 3.2* |
| **HexNAc(3)Hex(6)** | | 0.2 ± 0.4 | 3.8 ± 0.7* | 3.3 ± 0.7* |
| **HexNAc(3)Hex(6)Fuc(1)** | 0.4 ± 0.3 | 0.6 ± 0.5 | 4.2 ± 1.9* | 3.9 ± 0.5* |
| **HexNAc(4)Hex(5)Fuc(1)** | | | 6.4 ± 2.2* | 6.4 ± 0.4* |
| **HexNAc(5)Hex(5)** | | 0.2 ± 0.3 | | |

**Supplementary Table S3. Glycoform abundance at N496 in rHuNCAM in untreated, EndoNF treated, EndoNF and mild acid hydrolysis treated, and mild acid hydrolysis treated samples. *, *p* < 0.05 compared to untreated.**

| | **Untreated** | **EndoNF** | **EndoNF and mild acid hydrolysis** | **Mild Acid Hydrolysis** |
|---|---|---|---|---|
| **Glycoforms** | **Mean ± SD** | **Mean ± SD** | **Mean ± SD** | **Mean ± SD** |
| **HexNAc(2)Hex(3)** | 0.7 ± 0.1 | 0.6 ± 0.1 | * | * |
| **HexNAc(2)Hex(4)** | 3.1 ± 0.7 | 3.6 ± 0.1 | 1.7 ± 0.2* | 1.9 ± 0.5* |
| **HexNAc(2)Hex(5)** | 86.5 ± 2.2 | 83.3 ± 2.3 | 49.3 ± 6.4* | 44.3 ± 0.7* |
| **HexNAc(3)Hex(3)** | 1.0 ± 0.1 | 1.0 ± 0.1 | * | * |
| **HexNAc(3)Hex(4)** | 0.2 ± 0.3 | 0.9 ± 0.1 | 5.2 ± 0.4* | 4.2 ± 0.1* |
| **HexNAc(3)Hex(4)Fuc(1)** | 4.2 ± 2.0 | 5.6 ± 0.1 | 1.5 ± 0.2 | 1.3 ± 0.02* |
| **HexNAc(3)Hex(5)** | | 0.6 ± 1.0 | 14.3 ± 6.8* | 7.4 ± 1.2* |
| **HexNAc(3)Hex(5)Fuc(1)** | 3.9 ± 2.3 | 1.4 ± 0.1 | 0.4 ± 0.7 | 2.3 ± 0.1* |
| **HexNAc(3)Hex(5)NeuAc(1)** | 0.4 ± 0.6 | 2.0 ± 0.2 | | * |
| **HexNAc(3)Hex(6)** | | 1.0 ± 1.2 | 10.6 ± 5.5* | 7.9 ±0.1* |
| **HexNAc(3)Hex(6)Fuc(1)** | | 0.2 ± 0.4 | | |
| **HexNAc(4)Hex(5)** | | | 17.0 ± 4.3* | 30 ± 0.5* |
| **HexNAc(4)Hex(5)Fuc(1)** | | | | 0.8 ± 0.1* |

**Supplementary Table S4. Glycoform abundance at N176 in IgG2 in untreated, EndoNF only treated, and EndoNF and mild acid hydrolysis treated samples. *, $p < 0.05$ between untreated versus EndoNF and mild acid hydrolysis treated.**

| | Untreated | EndoNF | EndoNF and mild acid hydrolysis |
|---|---|---|---|
| **Glycoforms** | **Mean ± SD** | **Mean ± SD** | **Mean ± SD** |
| **HexNAc(3)Hex(3)** | | 0.02 ± 0.03 | |
| **HexNAc(3)Hex(3)Fuc(1)** | 3.2 ± 0.4 | 3.0 ± 0.5 | 2.5 ± 1.2 |
| **HexNAc(3)Hex(4)Fuc(1)** | 0.3 ± 0.4 | 0.6 ± 0.3 | 0.6 ± 0.6 |
| **HexNAc(3)Hex(4)Fuc(1)NeuAc(1)** | | 0.03 ± 0.1 | |
| **HexNAc(4)Hex(3)** | 0.2 ± 0.04 | 0.1 ± 0.1 | 0.5 ± 0.02* |
| **HexNAc(4)Hex(3)Fuc(1)** | 18.6 ± 9.5 | 10.7 ± 2.4 | 12.3 ± 1.9 |
| **HexNAc(4)Hex(4)** | | | 0.3 ± 0.5 |
| **HexNAc(4)Hex(4)Fuc(1)** | 31.0 ± 5.4 | 35.4 ± 0.9 | 37.8 ± 1.2 |
| **HexNAc(4)Hex(4)Fuc(1)NeuAc(1)** | 6.9 ± 1.3 | 7.3 ± 0.9 | * |
| **HexNAc(4)Hex(5)** | 0.1 ± 0.1 | 0.2 ± 0.2 | 0.5 ± 0.03* |
| **HexNAc(4)Hex(5)Fuc(1)** | 23.5 ± 2.5 | 23.8 ± 2.0 | 37.5 ± 1.3* |
| **HexNAc(4)Hex(5)Fuc(1)NeuAc(1)** | 7.9 ± 1.5 | 9.1 ± 0.2 | * |
| **HexNAc(4)Hex(5)NeuAc(1)** | | 0.02 ± 0.04 | |
| **HexNAc(5)Hex(3)Fuc(1)** | 4.4 ± 2.6 | 3.2 ± 3.3 | 4.5 ± 2.9 |
| **HexNAc(5)Hex(4)Fuc(1)** | 2.6 ± 2.3 | 5.0 ± 0.5 | 3.1 ± 2.4 |
| **HexNAc(5)Hex(4)Fuc(1)NeuAc(1)** | | 0.04 ± 0.1 | |
| **HexNAc(5)Hex(5)Fuc(1)** | 1.2 ± 0.2 | 1.4 ± 0.04 | 0.5 ± 0.8 |
| **HexNAc(5)Hex(5)Fuc(1)NeuAc(1)** | | 0.1 ± 0.1 | |
